# Visual landmarks calibrate auditory space across eye movements

**DOI:** 10.1101/853739

**Authors:** David Aagten-Murphy, Martin Szinte, Robert Taylor, Heiner Deubel

## Abstract

Visual objects that are present both before and after eye movements can act as landmarks, aiding localization of other visual stimuli. We investigated whether visual landmarks would also influence auditory localization – despite participants’ head position remaining unchanged. Participants made eye-movements from central fixation to a peripheral visual landmark, which either remained stationary or was covertly displaced. Following the movement, participants judged whether a stimulus (auditory or visual) was shifted in location relative to before the movement. Visual localization estimates shifted along with the landmark, although the landmark displacement itself went unnoticed. Interestingly, auditory localization estimates were also displaced. Thus, despite identical auditory input reaching the ears, two auditory stimuli originating from the same position were perceived as spatially distinct when the visual landmark moved. These results are consistent with the idea that auditory spatial information is encoded within an eye-centered reference frame and subject to spatial recalibration by visual landmarks.

**Highlights:**

- Visual landmarks affect stimulus localization across eye movements
- We show this also for auditory stimuli, even when the head remains stable
- Due to a visual landmark displacement, identical auditory stimuli are perceived as shifted
- This suggests that auditory space is calibrated on eye-centered maps across saccades

## Introduction

When locating objects in our environment we often have access to spatial information from multiple senses. However, because different modalities are encoded by different sensory bodies, each sensory input is encoded within a distinct reference frame. While visual information is encoded relative to the location of our gaze, auditory information is encoded relative to the orientation of our head (King, 2009). Integrating information from the same spatial location in these different sensory modalities is essential to our ability to identify and respond to multisensory objects. However, this integration requires that different sensory inputs can be precisely spatially aligned. As we can move our head, eyes and body position independently, this presents a considerable challenge. Indeed, every eye-movement causes substantial shifts of visual input projections on the retina but leave auditory input unchanged. For example, when making a large saccade—usually accompanied by a small head movement—accurate localization after the saccade requires that we know how far the head and the eyes have moved both relative to each other and to the external world (Lee and Groh, 2012). How does our brain maintain such a stable multi-sensory percept across saccades?

Internal signals estimating the extent of our movement provide one critical source of information. Experimental evidence has shown that efference copy, a copy of the motor command signaling an upcoming movement, is sent to perceptual areas (Wurtz, 2008) and plays an important role by signaling the anticipated magnitude of an upcoming saccade (Collins, 2010; Sommer and Wurtz, 2002). This pre-saccadic estimate of the saccade alter visual receptive fields (Duhamel et al., 1992) and is believed to be involved in our impression of a stable world across saccades (Cavanagh et al., 2010; Hall and Colby, 2011; Rolfs and Szinte, 2016). However, the necessity of an active prediction in generating perceptual stability remains controversial (Higgins and Rayner, 2015) amid suggestions that stability may simply be the default state of the perceptual system unless there is (considerable) evidence to the contrary. After the saccade, additional proprioceptive feedback from the eye muscles is available, which may provide an additional cue to the saccade magnitude (Poletti et al., 2013; Wurtz, 2008). However, studies have suggested its involvement in maintaining stable spatial percepts is limited, arguably due to their imprecision, and likely only situational (Higgins and Rayner, 2015; Wang et al., 2007). While these different extra-retinal signals are noisy (Collins et al., 2009), even in the absence of other cues they appear to be sufficient to enable a visual objects seen before the saccade to be accurately localized afterward (Collins et al., 2009; Deubel et al., 1998). This requires that remembered spatial information about the pre-saccadic stimulus is updated accurately by the estimated saccade vector (Aagten-Murphy and Bays, 2018; Collins et al., 2009).

Additionally, comparing the remembered location of visual objects before a saccade to their location after may also be used to estimate how far the eyes previously moved. Indeed, if the saccade target remains visible after the movement, the accuracy of the saccade can be inferred from the proximity of the target to the fovea (Cassanello et al., 2016; Collins and Wallman, 2012). This influence is most apparent when – rather than remaining stationary throughout the movement – the saccade target is covertly displaced mid saccade. Not only is this displacement not perceived by participants, but the post-saccadic information about the saccade targets appears to dominate internal signals, resulting in secondary, corrective movements and localization estimates being also shifted (Bridgeman et al., 1975; Deubel et al., 1996; Li and Matin, 1990; McConkie and Currie, 1996; Niemeier et al., 2003). Under these circumstances, the saccade target appears as a single object remaining stationary in the external world - even though its location after the saccade is considerably changed. In turn, discrepancy with the internal estimates is attributed to error in those estimates, with consistent correction leading to saccadic adaptation (Aagten-Murphy and Bays, 2018; Collins and Wallman, 2012; Lappe et al., 2000; Zimmermann and Lappe, 2010). As objects in the real world are unlikely to suddenly move during a saccade (Deubel et al., 1998), the perceptual system weights this visual information highly. The spatial range in which individuals perceive the visual object as remaining stable, and in which displacements of the target will go unnoticed, thus represents the range in which visual information not only informs our perceptual system about the extent of movement but also dominates other extra-retinal cues. Interestingly, this range of tolerable displacements is orientated in the direction of the saccade and is roughly equivalent to typical saccadic variability (Wexler and Collins, 2014). This fits well with recent findings demonstrating a near-optimal integration of retinal and extra-retinal information, depending critically on the reliability of the respective signals (Ostendorf and Dolan, 2015).Overall it appears that our perceptual system considers the precision of ocular, motor and visual cues, with a high weighting for post-saccadic visual cues when available, to form the best possible estimate of the how far the eye has moved.

Even when other visual objects are not task-irrelevant, their presence may influence this post-saccadic re-orientation process. Indeed, when individuals are required to memorize the location of a briefly presented visual probe seen before an eye-movement in the presence of another visual stimulus (acting as a visual landmark), their localization after the saccade depends on the direction and magnitude that the landmark was displaced (Deubel, 2004; Deubel et al., 2010, 1998). This suggests that the location of the visual landmark after the saccade not only influences estimates of saccade magnitude, but also the remembered location of objects held in spatial memory. Faced with the visual landmark not appearing in the predicted location after the saccade, the perceptual system appears to weight the visible location of the landmark stronger than its internal estimates of the saccade. This updating not just of currently visible information, but also information held in spatial memory, would help stitch together our experiences from before and after the disruption of the saccade. When dealing with multiple sensory modalities, which can be independently shifted relative to each other, maintaining alignment both within-modalities and across- is critical. How then does the shifting of a visual landmark affect other sensory modalities, such as the localization of an auditory sound? Will auditory localisation estimates also be influenced by the landmark, despite auditory information being encoded within a separate head-centered reference frame? Or will the modalities remain independent, risking the introduction of misalignment between the senses?

We sought to address this question by investigating the ability of participants to discriminate changes in the spatial position of either auditory or visual stimuli presented across an intervening saccade. By manipulating the location of visual landmarks across eye movements (Deubel, 2004; Deubel et al., 2010, 1998), while asking observers to localize a stimulus presented before the saccade, we could investigate how vision influences auditory localization across saccades. Importantly, as auditory information is encoded relative to the head, there is no reason for shifts in eye-gaze to influence auditory inputs if the head remains still. Thus, regardless of how far the participant moves their eyes and whether post-saccadic visual information affects their estimate of saccade length, the localization of auditory stimuli should remain unaffected. However, as evidence exists that auditory spatial information may also be transformed into eye-centered co-coordinates (Groh et al., 2001; Jay and Sparks, 1987, 1984; King et al., 1988; Lee and Groh, 2012; Russo and Bruce, 1994; Stricanne et al., 1996) auditory localization may also be affected by displaced landmarks.

## Results

At the start of each trial both the *fixation target*, an illuminated LED in the center of the display, and the *visual landmark*, two vertically displaced illuminated LEDs in the periphery, were visible. In the saccade experiment participants initiated a saccade to the middle of the visual landmark when the fixation LED was extinguished. During this initial fixation period, an auditory or a visual stimulus (*pre-saccadic stimulus*) was presented in the periphery at one of five pre-saccadic stimulus locations (±9°, ±10.5°, ±12°, ±13.5°, ±15° relative to the central fixation). In the main experiment, participants were then required to maintain fixation until the fixation LED was extinguished, triggering the saccade. While the eyes were moving, the visual landmark could either remain stationary or be displaced by up to three visual degrees, inward or outward (Figure 1 and Experimental Procedure). Because the displacement of the visual landmark occurred while the eyes were moving, any changes in position went undetected due to saccadic suppression of displacement (Bridgeman et al., 1975). The second stimulus (*post-saccadic stimulus*) was then presented 500 ms after the initiation of the saccade and was always the same modality as the first. Finally, participants were required to judge whether this second stimulus had originated from the left or right of the initial pre-saccadic stimulus. The ability to spatially localize auditory and visual stimuli across a saccade was quantified by determining the accuracy (point of subjective equality; PSE) and precision (just-noticeable-difference; JND) with which discrepancies in the location of the pre- and post-saccadic stimuli were discriminated.

**Figure 1.**
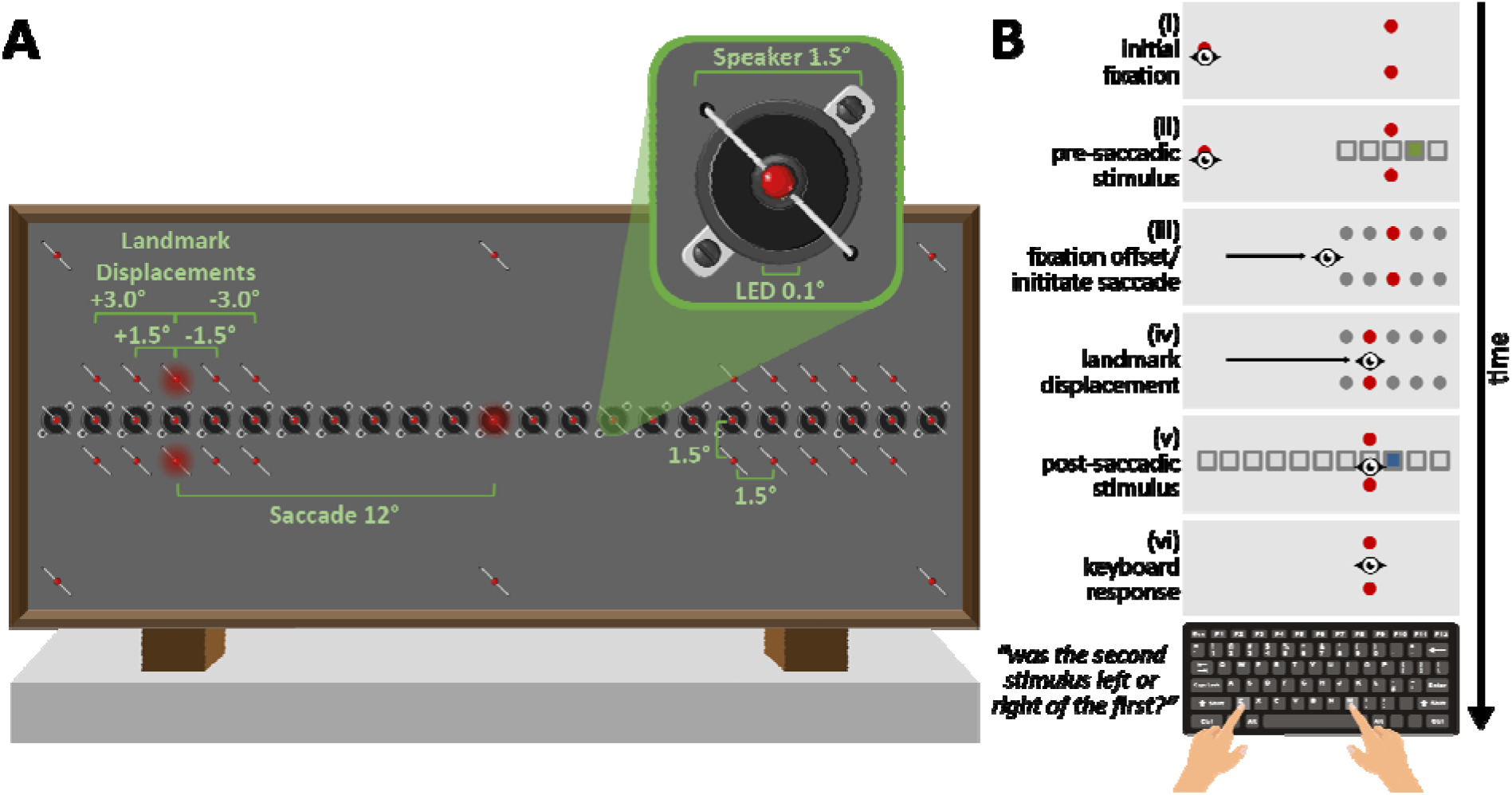
Experimental procedure. **A. Apparatus**. Custom setup consisting of 23 speakers and 43 LEDs (with an additional 6 for eye-tracking calibration). The setup was covered with a thin (not shown) auditory transparent material to prevent participants seeing the stimuli locations. **B. Trial sequence**. (i) Participants fixated on a central LED while a visual landmark, two vertically displaced LEDs, was presented in the periphery. (ii) A brief visual or auditory pre-saccadic stimulus (50 ms; green box) was next presented at one of five locations near the visual landmark (see grey boxes). (iii) Participants were instructed to move their eyes to the visual landmark at the offset of the fixation target. (iv) During the saccade, the visual landmark could either remain stationary or be displaced (see grey dots). (v) Shortly after the saccade a second visual or auditory stimulus (blue box) was played from one of twelve different locations (see grey boxes). (vi) Participants responded via the keyboard whether the second (post-saccadic) stimulus was located left or right of the first (pre-saccadic) stimulus.

### Auditory localization during fixation

Before performing the main saccade experiment all participants took part in an initial fixation experiment, stable fixation throughout the entire trial at either a central (0°) or eccentric (±12°) location (see Supplemental Information). The results of the initial fixation experiments demonstrated that participants could both accurately (less than 1° bias for both central and eccentric fixation positions) and precisely (about 3° JND) localize auditory stimuli when required to hold their gaze at either a central or a peripheral fixation location.

### Visual and auditory localization across a saccade with static landmarks

To examine the influence of the visual landmark on auditory and visual localization we contrasted trials in which the landmark remained stationary with those in which it was displaced during the saccade (Figure 2). For the trials in which the visual landmark remained at the same location (Figure 2A; middle row: “no shift”), visual stimuli were localized across a saccade without bias (mean ± SE; PSE = −0.21 ± 0.11°, t(7) = −1.92; *p* = 0.1) and with high precision (JND = 0.89 ± 0.07°), with the post-saccadic PSE changing reliably with the pre-saccadic location (SLOPE = 0.93 ± 0.13°; t(7) = 6.94; *p* < 0.01) and not differing from unity (t(7) = −0.56; *p* = .60). This indicates participants were able to accurately localize the horizontal location of the different pre-saccadic visual stimuli. Additionally, the precision of their localization judgements did not depend on pre-saccadic stimulus eccentricity (SLOPE = −0.03 ± 0.03; t(10.65) = −1.05; *p* = 0.32).

**Figure 2.**
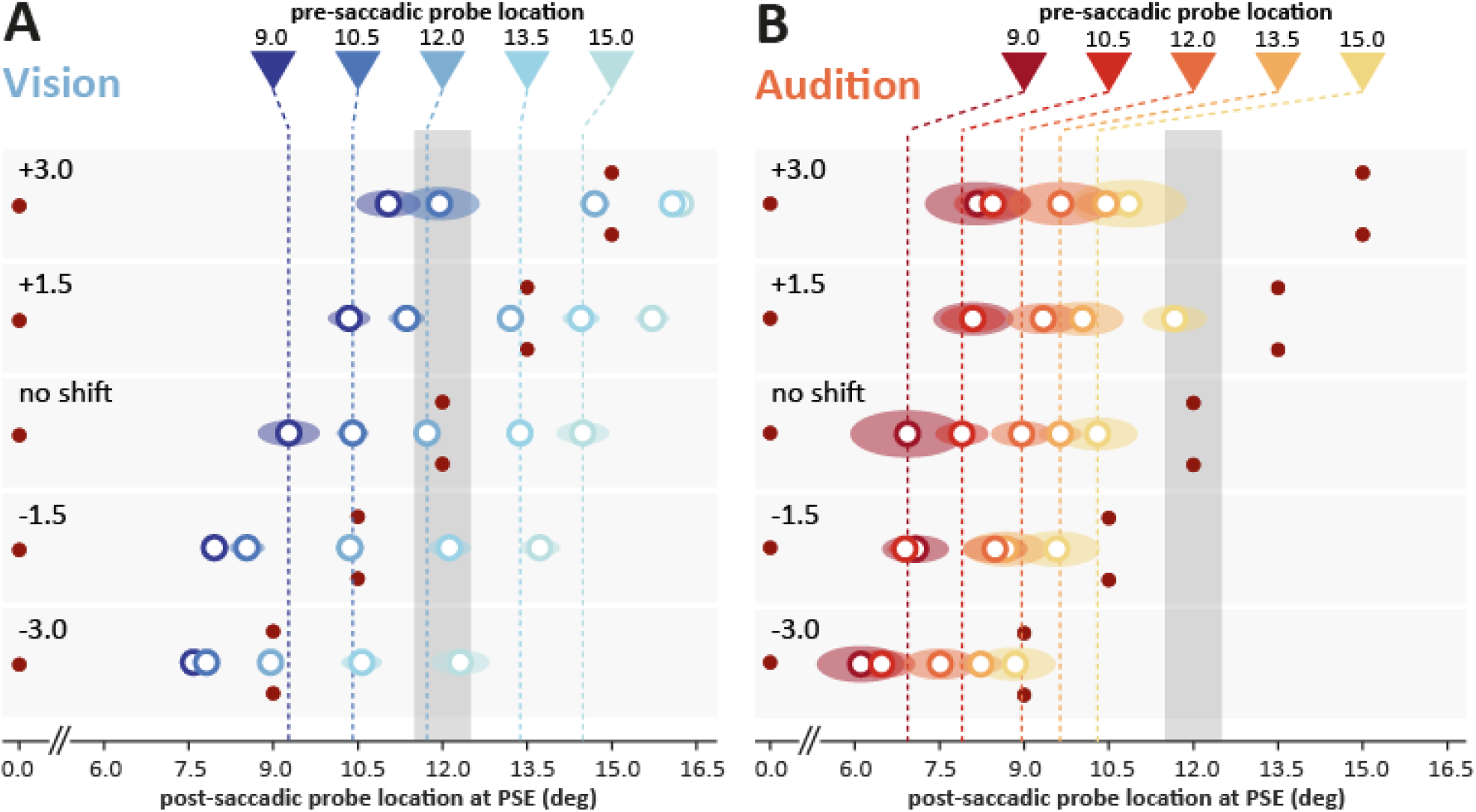
Auditory and visual localization. Averaged point of subjective equality (i.e. the location at which the post-saccadic stimulus was perceived as aligned to the pre-saccadic stimulus) for visual (**A**) and auditory stimuli (**B**), the shift of the visual landmarks (rows). The averaged location at which a post-saccadic stimulus was reported as occurring at the same location as the pre-saccadic stimulus (point-of-subjective equality; PSE) for the different pre-saccadic stimulus eccentricity are shown in different colors matching the corresponding colored arrows. The dark gray shaded region shows the pre-saccadic position of the visual landmark (at 12°). The vertical dashed lines indicate the PSE observed without a shift of the visual landmark and can be seen as a baseline. The shaded colored ovals represent the SEM.

In contrast, participants’ auditory localization responses exhibited a clear bias in the opposite direction of the saccades (PSE = −3.23 ± 0.35°, t(7) = −9.3; *p* < 0.01). Planned contrasts revealed that this inwards bias was larger than what was observed for audition during the central (diff = 3.40 ± 0.51°; t(7) = 6.71; *p* < 0.01) and peripheral fixation task (diff = 2.48 ± 0.31°; t(7) = 7.93; *p* < 0.01) and was the strongest for more peripheral pre-saccadic stimulus locations (presented from 9 to 15° from the fixation target: −2.00 ± 0.63°; −2.70 ± 0.47°; −3.00 ± 0.53°; −3.73 ± 0.61°; −4.73 ± 0.72°). While post-saccadic PSE reliably changed with the pre-saccadic location (SLOPE = 0.57 ± 0.11°; t(7) = 4.98; *p* < 0.01), the slope also significantly deviated from unity (t(7) = −3.77; *p* < 0.01). This indicates that the range of auditory responses across different pre-saccadic locations was compressed and exhibited a central tendency (i.e. a bias towards the average stimulus location; (Karaminis et al., 2016; Watson, 1957)).

Our results reveal that pre-saccadic auditory stimuli were on average mislocalized inwards by 26.9 ± 2.9% of the saccade size. This means that a post-saccadic auditory stimulus had to be presented approximately 3° closer to the fixation target (Figure 2B) to be perceived as originating from the same location as the pre-saccadic stimulus. As for vision, the precision of auditory responses (JND = 3.05 ± 0.20°) did not depend on the eccentricity of the pre-saccadic stimulus (SLOPE = −0.03 ± 0.05°; t(16.34) = −0.60; p = 0.56). Indeed, planned contrasts demonstrate that the precision of auditory localization across a saccade was not significantly different from either central (diff = 0.41 ± 0.16°; t(7) = 2.61; *p* = 0.11) or peripheral (diff = −0.05 ± 0.17°; t(7) = −0.29; *p* = 0.78) fixation. However, as could be expected when comparing across modalities (Alais and Burr, 2004), auditory responses were overall less precise than visual responses (diff = 2.16 ± 0.20°; t(7) = 10.83; *p* < 0.001).

### Visual and auditory localization across a saccade with displaced landmarks

For the trials in which the visual landmark was displaced during the saccade, we found different biases for the different sizes and directions of the visual landmark displacement (Figure 2 and Figure 3A) for both visual (landmark displaced by −3° to 3° during the saccade: −3.01 ± 0.10°; −1.57 ± 0.09°; −0.21 ± 0.11°; 1.00 ± 0.11°; 2.00 ± 0.20°) and auditory targets (−4.57 ± 0.31°; −3.86 ± 0.28°; −3.23 ± 0.35°; −2.601 ± 0.37°; −2.52 ± 0.53°). These estimates reveal that post-saccadic localization for both sensory modalities exhibited a systematic mislocalization depending on the amount of visual landmark displacement.

**Figure 3.**
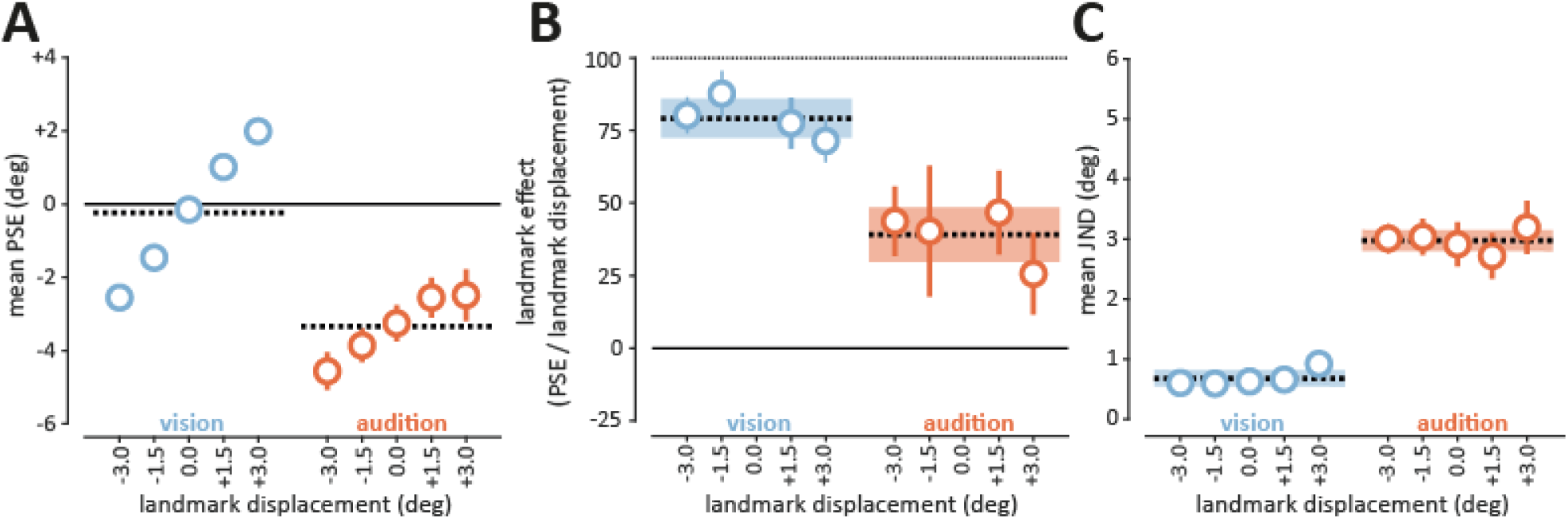
Accuracy, visual landmark effect and precision of the visual and auditory stimuli localization. **A.** Averaged localization accuracy (PSE) observed as a function of the visual landmark displacement. **B.** Averaged proportion of the visual landmark effect (proportion of the stimulus mislocalization relative to the visual landmark displacement). **C.** Averaged localization precision (JND) observed as a function of the visual landmark displacement. Data are shown for visual (blue) and auditory (orange) stimuli. Error bars show SEM. Dashed lines show the average across the different visual landmark displacement, and the colored shaded regions show SEM.

To better quantify the localization changes we fit the data with linear mixed models enabling us to separately estimate any biases common to all displacement conditions, how localization estimates changed for different pre-saccadic stimulus locations, and the effect of displacing the visual landmark. With the increased power of examining all displacement conditions, we found that visual estimates across a saccade were biased slightly (but consistently) inwards (−0.35 ± 0.07°; t(7.72) = −4.89; *p* = 0.001), while auditory estimates were again found to be considerably biased towards fixation after an intervening saccade (−3.36 ± 0.33°; t(7.00) = −10.03; *p* < 0.001). Although localization estimates varied with pre-saccadic location for both modalities (vision: t(7.01) = 9.73; *p* < 0.001; audition: t(7.00) = 6.35; *p* < 0.001), there was a considerable compression of the response range for auditory localization (0.51 ± 0.08), while visual localization remained veridical with pre-saccadic location (1.00 ± 0.1).

Critically, both visual and auditory localization were found to depend on the degree of visual landmark displacement. Visual localization shifted by 82.8% ± 3.6% of the displacement of the visual landmark (t(7.32) = 23.05; *p* < 0.001), and auditory localization shifted by 35.7 ± 5.2% (t(7.10) = 6.870; *p* < 0.001). This reveals that not only is there a consistent inwards bias in auditory localization across a saccade, but there is an independent, additional shift in localization depending on the post-saccadic location of the visual landmark.

While visual precision (0.91 ± 0.08°) did not change across different pre-saccadic stimulus locations (t(12.88) = 1.63; *p* = 0.13), it did vary with landmark displacement (t(14.95) = 3.78; *p* = 0.002) such that JND changed by 6.3 ± 1.7% of the landmark displacement. However, a significant interaction between pre-saccadic stimulus location and landmark displacement (t(36.74 = −4.89; *p* < 0.001)—and inspection of the data—revealed that this effect was driven by the most extreme pre-saccadic location reaching the edge of the testable range and not by a meaningful effect of landmark displacement on the precision of visual localization.

Importantly, despite considerable changes in auditory localization, the precision of auditory estimates did not change with the degree of landmark displacement. Indeed, auditory JND remained at 3.11 ± 0.17° for all pre-saccadic locations (t(9.91) = −0.07; *p* = 0.95) and for all displacements (t(9.05) = 0.93; *p* = 0.38). Thus, the considerable effect of shifting visual landmarks on auditory localization did not appear to increase the uncertainty involved in localization, but instead represented a rigid displacement in position without changes in precision.

## Discussion

We investigated the influence of visual landmarks on the localization of auditory stimuli across saccades. To do so, participants were required to remember the location from which a pre-saccadic visual or auditory stimulus occurred and compare it to the location of a post-saccadic stimulus of the same sensory modality. We systematically manipulated whether the visual landmark was displaced during the saccade, when the movement of the eyes prevented its detection (Bridgeman et al., 1975). Our results revealed that the memorized location of the pre-saccadic visual stimulus was shifted on average by 79% of the trans-saccadic displacement of the visual landmark. This supports previous work showing the influence of visual landmarks on the localization of other visual stimuli (Deubel, 2004; Deubel et al., 2010, 1998). Critically, we also examined how auditory localization was influenced by a visual landmark. We found that memorized auditory stimuli were shifted on average by 39% of the trans-saccadic visual landmark displacement. Thus, the post-saccadic visual landmark exerted a considerable influence on the localization of remembered stimuli, despite remembering the landmark itself not being required to perform the task. Importantly, though the visual landmark induced substantial shifts in the localization of both modalities, there were no changes in localization precision as a function of the visual landmark displacement. This indicates that the landmark-induced shifts did not arise due to increased spatial uncertainty. Instead, our results suggest that the displacement of the visual landmark caused the spatial memory for the pre-saccadic stimulus to undergo a rigid, albeit incomplete, translation in space. This surprising influence of visual landmarks on the localization of auditory stimuli suggests that vision is used to calibrate auditory spatial representations across the saccade.

While spatial visual information is based on the direct projection from an object onto the retina (eye-centered reference frame), spatial auditory information of an object has to be indirectly extracted from the sound spectral cues, timing and intensity differences between the two ears (head-centered reference frame, (Keating and King, 2015; King et al., 2001)). This means that accurate audiovisual spatial localization relies upon two different reference frames being spatially aligned. The precision of auditory spatial information is typically inferior to visual spatial information, accordingly this calibration has been thought to take place primarily via visual space (King, 2009; Lee and Groh, 2012; Recanzone, 2009; Wozny and Shams, 2011). Indeed, modulation of auditory space by eye position has been found in multiple cortical and sub-cortical regions in both human and animal studies, including the Frontal Eye Fields (Russo and Bruce, 1994), the Superior Colliculus (Campos et al., 2006; Hartline et al., 1995; Jay and Sparks, 1987, 1984; Opstal et al., 1995; Paré and Munoz, 2001; Peck et al., 1995; Populin et al., 2004; Zella et al., 2001), the Inferior Colliculus (Groh et al., 2001; Zwiers et al., 2004), as well as various sites within the intra-parietal cortex (Mullette-Gillman et al., 2009, 2005; Schlack et al., 2005; Stricanne et al., 1996). Recently, ocular signals related to saccade direction and magnitude have even been found to alter eardrum behavior (Gruters et al., 2018), suggesting spatial interactions between the senses to occur from even the earliest levels of auditory processing. While it was traditionally thought that sensory information was stored in a singular reference frame (Batista et al., 1999; Cohen and Andersen, 2002), recent studies provide strong evidence that it is instead stored in parallel within multiple different reference frames (see (Heed et al., 2015)). However, how spatial information from different sensory modalities comes together to provide us with a singular percept remains an open question.

Our results revealed a modification in auditory localization based on both the eye position (Goossens and Opstal, 1999; Klingenhoefer and Bremmer, 2009; Krüger et al., 2016; Pavani et al., 2008; Vliegen et al., 2004) and the location of visual references. This suggests that not only are auditory targets influenced by the position of the eye but that they are also partially encoded into an eye-centered reference frame. We suggest that it was the modulation of this eye-centered reference frame by the displacement of the visual landmark that induced the systematic shifts in auditory localization observed. Typically, information about the extent of an upcoming saccade (efference copy or corollary discharge; (Bridgeman, 2007; Sommer and Wurtz, 2002)) is transmitted to retinotopic visual areas to allow the remapping of visual space (Duhamel et al., 1992). Such a mechanism could then facilitate object correspondence despite drastic changes in the object projection on the retina. This compensation is considered essential in allowing the detection of displacements across saccades (Boon et al., 2016; Bridgeman et al., 1975; Deubel et al., 1996), integrating visual information pertaining to the same object (Aagten-Murphy and Bays, 2018; Ganmor et al., 2015; Szinte et al., 2016; Wijdenes et al., 2015; Wolf and Schütz, 2015), and providing perceptual stability (Cavanaugh et al., 2016; Rolfs, 2015; Rolfs and Szinte, 2016). Upon landing, the perceptual system has access to motor information about the magnitude of the intended movement, proprioceptive information about the movement executed, and a visual error signal indicating the difference between the actual and predicted post-saccadic visual input (Atsma et al., 2016; Collins et al., 2009; Niemeier et al., 2003; Ostendorf and Dolan, 2015). By integrating these different signals, the perceptual system can ensure both that our spatial representations remain stable across eye movements and that our eye movements remain accurate.

We propose that the shift of the visual landmark during the saccade induced an error signal upon saccade landing, as the visible location of the landmark after the saccade was not at the predicted location given the intended eye movement. To enable stimuli seen before the saccade to be localized afterwards, their eye-centered representation must then similarly be remapped by the size of the saccade in its opposite direction (Szinte et al., 2018).

This means that the perceptual system will anticipate that the visual landmark (here also the saccade target) would, after the correctly executed eye movement, be subsequently located at the fovea. However, when the visual landmark is displaced during the saccade, although the efference copy would signal that the eyes had correctly moved, the post-saccadic retinal input would be inconsistent with this action. This creates a conflict between different sensory cues coding for the magnitude of the saccade, with the discrepancy consistent with either a shift in the location of the visual landmark, or with an unexpected error in the execution of the movement. Due to the strong assumption of visual stability (Atsma et al., 2016; Deubel et al., 1998; Niemeier et al., 2003), related to the unlikelihood of an otherwise stationary visual landmark suddenly and abruptly changing position precisely during the saccade, the majority of this error should be attributed to an inaccuracy in the movement. Therefore, to ensure that spatial representations will be correctly aligned to the external world, sensory representations must be further shifted by the same magnitude as the visual landmark displacement to account for the (seemingly) inaccurate initial estimate of the saccade magnitude. The result of this is that the remembered location of visual targets presented before the saccade will also be shifted, based on the degree of displacement of the visual landmark.

However, for a similar process to influence auditory localization, this would require that auditory information was also encoded within an eye-centered reference frame. While it is important to note that the landmark effects for auditory stimuli were substantially smaller than for visual stimuli, the reduced effect could be consistent with participants’ final localization responses for auditory stimuli representing the integration of shifted eye-centered auditory information and residual, unchanged head-centered auditory information.

This interpretation of the influence of a visual landmark is consistent with other studies in which the accuracy of eye or hand movements towards remembered visual targets was found to be affected by the presence of a displaced landmark (Byrne and Crawford, 2010; Camors et al., 2015; Deubel, 2004; Fiehler et al., 2014; Li et al., 2017; Schütz et al., 2013). Indeed, supporting the idea that a strong assumption for visual stability acts as a prior for landmark stability, studies in which the reliability of the visual landmark was explicitly reduced (by jittering its spatial location) demonstrated that participants’ localization responses became noticeably less reliant on the landmark’s location (Byrne and Crawford, 2010). Similarly, studies of saccadic suppression of displacement have shown that when the displacement of the saccade target is too large (such that such a large eye error is highly unlikely) the assumption that changes are due to errors in the eye movement is broken (Atsma et al., 2016). However, the generalization of these effects to the auditory modality is remarkable and underlines the importance of visual landmarks in also maintaining alignment between the senses across movements.

It is important to distinguish these landmark shifts from other spatial effects such as audiovisual capture and peri-saccadic mislocalization. In audiovisual capture, the onset of a visual stimulus causes a simultaneously presented auditory stimulus to be perceived as originating from the visual location (Alais and Burr, 2004). If the visual landmark in our experiment had captured the auditory stimulus, then localization responses would converge towards it. Instead, for each of the different landmark displacements, auditory localization responses were shifted consistently in the same direction. Additionally, to reduce the likelihood of audiovisual capture, the onset of the pre- and post-saccadic visual landmark were separated by at least 450 ms from the appearance of the auditory stimuli. This exceeded the typical range in which capture effects are observed (Lewald and Guski, 2003). In the peri-saccadic mislocalization of sounds (Klingenhoefer and Bremmer, 2009), a 5 ms auditory stimulus occurring in the temporal vicinity of the saccade was perceived as shifted. However, when stimuli occurred more than 200 ms before or after the saccade no shift was observed. As our auditory stimuli were also played more than 450 ms from the saccade, we can be confident that our landmark shifts do not arise from peri-saccadic mislocalization.

In addition to landmark-induced localization shifts, we also observed a general modulation of auditory localization by the execution of an eye movement, in the absence of any displacement of the visual landmark. This resulted in a systematic shift of the remembered position of auditory targets in the opposite direction of the saccade and suggests that, although the perceptual system can accurately account for the intervening saccade during the localization of visual stimuli (Szinte and Cavanagh, 2011), the execution of the eye-movement introduced biases into auditory localization. Previously, some auditory localization studies have reported little or no influence of an eye movement (Goossens and Opstal, 1999; Klingenhoefer and Bremmer, 2009; Vliegen et al., 2004), others have shown substantial shifts in the opposite direction to the saccade (Krüger et al., 2016; Pavani et al., 2008), suggesting that the existence of these eye-related auditory biases is highly sensitive to the specific experimental paradigm utilized. In particular, aspects of the design might influence an individual’s priors at the decision-making stage (Grüsser, 1983; Lewald and Ehrenstein, 1998; Parise et al., 2011).

Our results suggest that visual landmarks are used to calibrate not only visual but also auditory space across eye movements. The importance of vision for auditory space has been shown in both animal and human studies. Raising owls with prismatic glasses leads to them developing systematic distortions in their auditory localization (Knudsen and Knudsen, 1985), while depriving ferrets of all visual signals results in them developing highly disordered auditory spatial maps (King and Carlile, 1993). Human studies using magnifying lenses have also shown systematic shifts in auditory spatial localization (Zwiers et al., 2003). Furthermore, the single presentation of a misaligned audio and visual stimulus, perceived as originating from the same position, is sufficient to temporarily alter where subsequently presented unimodal auditory stimuli are localized (Wozny and Shams, 2011). These studies, using visual adaptation methods, demonstrate the susceptibility of auditory space to a calibration by vision (King, 2009; Recanzone, 2009; Wozny and Shams, 2011). In our results, without relying upon adaptation, we showed that the retinal error of a task-irrelevant visual landmark was able to shift auditory spatial representations on a trial-by-trial basis. While such a mechanism may appear to result in a decrease in the accuracy of spatial localization across a saccade, in typical viewing scenes in which the stable visual landmarks are indeed a reliable cue to space in the external world, this instead assists in the maintenance of a stable, multi-sensory perception of the surroundings.

## Experimental procedures

### Participants

Eight students or staff of the LMU München (including one author) participated in the experiments. All participants had normal or corrected-to-normal vision, were naive as to the purpose of the study and received compensation of 10 Euros per hour of testing. The experiments were undertaken with the understanding and written consent of all participants and were carried out in accordance with the Declaration of Helsinki and the requirements of local ethics review board at LMU München for experiments involving eye tracking.

#### Setup

Participants sat in a dark, sound-attenuated cabin, with their head both held steady with a chin and forehead rest across all trials. The cabin’s walls, floor, ceiling, and all room furniture were covered with black sound-absorbing foam to eliminate acoustic reflections. The experiment was controlled by a Hewlett-Packard Intel Core i7 PC (Palo Alto, CA, USA) located outside the cabin. Manual responses were recorded via a standard keyboard. The dominant eye’s gaze position was recorded and available online using an EyeLink 1000 Desktop Mounted (SR Research, Osgoode, Ontario, Canada) at a sampling rate of 1000 Hz and equipped with an invisible 940 nm infrared illuminator to prevent the eye tracker to act as a visual reference. The experimental software controlling the display, the response collection as well as the eye tracking was implemented in Matlab (MathWorks, Natick, MA, USA), using the Psychophysics and EyeLink toolboxes (Brainard, 1997; Cornelissen et al., 2002; Pelli, 1997). Stimuli were presented at a viewing distance of 114 cm on a custom-made audiovisual screen. Auditory signals were played from 23 speakers (0.75° radius, auditory stimuli) arranged on the horizontal meridian of the screen and located to the right and left side (at eccentricities of ±1.5°, ±3°, ±4.5°, ±6°, ±7.5°, ±9°, ±10.5°, ±12°, ±13.5°, ±15°, ±16.5°) of a central speaker (0°). Sounds were played via a 24 Input/Output sound card (MOTU, Cambridge, MA, USA) amplified by two PowerPlay Pro amplifiers (Behringer, Wellich, Germany). Visual signals were displayed by 49 red-LEDs (0.05° radius) which were controlled at 1000 Hz via an Arduino Due electronic board (Arduino, Turin, Italy). 23 of the LEDs (*visual stimuli*) were mounted on top of the 23 speakers and an additional 20 LEDs (*visual landmark*) mounted 1.5° above and below the eccentric landmark locations (at eccentricities of ±9°, ±10.5°, ±12°, ±13.5°, ±15°). The remaining 6 LEDs were mounted on the screen corners and at the top and bottom of the screen vertical meridian to facilitate the calibration of the eye tracker using a custom 9-point calibration pattern. A sound-transparent spandex screen covered the entire display to ensure that participants remained unaware of the hardware layout.

### Procedure

Once subjects were comfortably seated in the soundproof cabin (see Setup), each trial began with an initial fixation period of 500 ms during which the participant’s gaze must remain within a 1.5° radius of the fixation target (central red LED) after which the visual landmark appeared. The visual landmark was composed of two vertically aligned LEDs (separated by 3°) located 12 degrees to the right or to the left of the screen center. Once the visual landmark had remained visible for 1000 ms the fixation target disappeared, indicating participants should saccade towards its center as quickly and accurately as possible (mean saccade latency: 226 ± 37 ms). Across trials, the visual landmark either stayed at the same position (20% of the trials) or was shifted as soon as a saccade was detected (80% of the trials) by either 1.5° or 3.0° inward or outward relative to the *fixation target* location. Saccades were detected online as soon as the eye position crossed a 1.5°-radius virtual circle centered on the *fixation target* (14 ± 2 ms after saccade onset). Both before and after the saccade brief auditory or visual stimuli were presented. The first stimulus occurred 500 ms after the appearance of the visual landmark (~726 ms before saccade onset) at one of 5 positions surrounding the visual landmark (±9°, ±10.5°, ±12°, ±13.5°, ±15 relative to the screen center; see supplementary details). The second stimulus occurred 500 ms after the online detection of the saccade and was played at one of the 11 positions on the same side as the landmark (between 0° and 16.5° in steps of 1.5°) for the auditory condition. For the visual condition, due to the higher precision of localization for vision, the range was reduced to better focus trials around the relevant portion of the display (between 7.5° and 16.5° in steps of 1.5°). The average inter-stimulus duration interval between stimuli for both modalities was 1264 ± 76 ms and pre- and post-saccadic stimuli were both separated by at least 450 ms from the saccade. The pre- and post-saccadic stimuli could either be a visual (50 ms illumination of a red LED) or auditory (50 ms broadband Gaussian white noise with a 5 ms raised cosine ramp for onset and offset). At the end of each trial, participants reported whether the second stimulus was played to the left or to the right of the first stimulus via the keyboard.

Participants completed approximately 1500 trials with a 12° saccade in the auditory condition and 1000 trials with a 12° saccade in the visual condition, separated into blocks of 100 trials. In each block trials with fixation breaks or incorrect saccades were immediately discarded and repeated at the end of each block in a random order. Participants performed these blocks over 2 to 3 experimental sessions of approximately 120 minutes (including breaks). Further offline analysis of eye movements resulted in the exclusion of 12% of trials in saccade conditions due to blinks, fixation errors, or incorrect saccades.

Before performing the saccade task, participants took part in an additional auditory localization at fixation task. This ensured all participants could reliably localize the auditory stimuli. The procedure was identical to that of the saccade task (outlined above) except that participants were required to maintain their fixation throughout the experiment and the visual landmark was never shifted (see Supplementary Material). For each fixation condition participants performed 10 blocks of 100 trials (with trials repeated when online errors were detected repeated) over 2 to 3 experimental sessions of 120 minutes (including breaks) for a total of 1000 trials in each condition. All participants completed the fixation task before starting the saccade task. Additional offline analysis of eye movements resulted in the exclusion of 2% of fixation trials due to blinks, or to fixation errors not detected online.

#### Data pre-processing

Eye-position data was further processed offline. Saccades were detected based on their velocity distribution (Engbert and Mergenthaler, 2006) using a moving average over twenty subsequent eye position samples. Saccade onset was detected when the velocity exceeded the median of the moving average by 3 SDs for at least 20 ms. We included trials if a correct fixation was maintained within a 1.5° radius centered on *fixation target* (all tasks), if a correct saccade started at *fixation target* and landed within a 2.5° radius centered on *visual landmark* (saccade tasks only), and if no blink occurred during the trial (all tasks).

### Behavioral data analysis

For each individual participant, and condition, we computed the accuracy and precision with which changes in the location of the pre- and post-saccadic stimuli could be discriminated. Accordingly, the parameters of interest were the point of subjective quality (PSE)—the location at which the second stimulus was equally likely to be reported as originating either to the left or right of the first stimulus—and the just noticeable difference (JND), which indexes of the precision of stimulus localization. To do this we estimated participant-level parameters using a generalized linear mixed-effects model (GLMM) with a probit link function. Specifically, for the *j*^*th*^ participant in the *k*^*th*^ condition, the proportion of observed responses, *r*_*ijk*_, to the *i*^*th*^ stimulus, *s*_*i*_, is modelled by

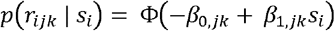

where *β*_0,*jk*_ ~ *N*(*B*_0,*k*_, *σ*_0,*k*_) and *β*_1,*jk*_ ~ *N*(*B*_1,*k*_, *σ*_1,*k*_), and Φ(·) is the Gaussian cumulative distribution function. The model accordingly assumes that the participant-level estimates (i.e., *β*_0,*jk*_ and *β*_1,*jk*_) vary randomly within each condition (i.e., a random intercept and slope model), and that this variation is best modelled by a Gaussian distribution centered on the group-level estimates (i.e., *B*_0,*k*_ and *B*_1,*k*_). This hierarchical structure constrains the dispersion in the participant-level estimates by shrinking larger estimates towards the group mean, thus mitigating the effect of extreme data points on measures of central tendency. Estimation of subject-level parameters were conducted using the lme4 package (Bates et al., 2015) in R (R Core Team, 2018). Additionally, we determined the strength of the landmark effect as the ratio of the localization bias to the magnitude of the visual landmark displacement.

For each experiment, all statistical tests were performed on the participant-level estimates. To examine whether there were any reliable changes in performance across conditions we used linear mixed-effects models (LMM) which were implemented using the afex package (Singmann et al., 2016) in R (R Core Team, 2018). All analyses assumed a full random-effects structure that included both by-participant random intercepts and slopes. However, owing to the random effects structure, conventional statistical tests on the fixed-effect coefficients cannot be straightforwardly applied. This is because the null distribution for the model parameters and test statistics are typically unknown, and thus require an approximation, or simulation-based approach, to test model effects. Accordingly, all inferential tests applied Satterthwaite’s method to approximate the degrees of freedom for both *t* and *F* tests, which can result in non-integer degrees of freedom. The method calculates the effective, or pooled, degrees of freedom (which can often return non-integer degrees of freedom) and have been shown to provide more robust *p*-values than likelihood-ratio tests (Kuznetsova et al., 2017). Excerpts of the key analysis performed are shown in the supplementary tables (Table S1–S6).

## Acknowledgements

This research was supported by a DFG grant (DE336/3-1) to HD and to M.S. (SZ343/1) and a Marie Sklodowska-Curie Action Individual Fellowship to M.S. (704537).

## Authors’ contributions

Conceptualization: D.A-M., M.S. and H.D.; Methodology: D.A-M., M.S. and H.D; Software: D.A-M. and M.S.; Formal analysis: D.A-M., R.T.; Investigation: D.A-M.; Writing—Original draft: D.A-M.; Writing—Review and editing: D.A-M., M.S., R.T. and H.D; Visualization: D.A-M.; Supervision: H.D.; Project administration: D.A-M., M.S. and H.D; Funding acquisition: M.S. and H.D..

## Supplementary Material

### Auditory localization at fixation

Participants initially performed a fixation version of the auditory localization task in which they maintained fixation throughout the entire trial at either the central fixation target (0°; *central fixation*) or at the location of the saccade target (±12°; *eccentric fixation*). During central fixation, when head and eye-gaze were aligned, participants localized auditory stimuli both accurately (BIAS = 0.17 ± 0.24°; t(7) = 0.69; *p* = 0.51) and precisely (JND = 3.46± 0.18°). Participants post-saccadic PSE reliably changed with the pre-saccadic location (SLOPE = 0.86 ± 0.03°; t(31.54) = 25.17; *p* < 0.01), although the slope deviated from unity (t(31.54) = −4.1; *p* < 0.01), indicating a slight range compression (i.e. regression towards the mean).

In contrast, when the head and eye-gaze were misaligned during eccentric fixation, participants localized auditory stimuli with a slight bias towards the central heading direction (BIAS = −0.75 ± 0.21°; t(7.02) = −3.64; *p* = 0.01). This effect was found to be predominantly driven by the most eccentric first stimulus locations (from 7.5 to 16.5°: −0.50 ± 0.21°; −0.13 ± 0.31°; −0.80 ± 0.33°; −0.47 ± 0.35°; −1.15 ± 0.37°; −1.32 ± 0.41°; −1.91 ± 0.44°). As with central fixation, post-saccadic PSE reliably changed with the pre-saccadic location (SLOPE = 0.76 ± 0.04°; t(7.67) = 17.08; *p* < 0.01) and showed a slight range compression (t(7.67) = −5.32; *p* < 0.01).

Additionally, participants’ auditory localization precision did not significantly differ (diff = 0.46 ± 0.20; t(7) = 2.54; *p* = 0.12) between peripheral (JND = 3.00 ± 0.23°) and central (3.46 ±0.18°) fixation conditions. Furthermore, for both central (SLOPE = − 0.01 ± 0.04; t(7) = 0.2; *p* = 0.85) and peripheral (SLOPE = − 0.01 ± 0.04; t(7) = −0.2; *p* = 0.85) fixation, there was no variance in JND across pre-saccadic standard locations.

Several behavioral studies have also shown that static discrepancies between eye-head reference frames can result in localization shifts for both visual (Harris and Smith, 2008; Kopinska and Harris, 2003; Lewald and Ehrenstein, 1998; Rossetti et al., 1994; Wexler, 2003) and auditory stimuli (Lewald and Ehrenstein, 1998, 1996; Weerts and Thurlow, 1971). In general, these studies have suggested that a discrepant eye-head alignment induces small (~3°) localization biases in the opposite direction of eye-gaze (Pavani et al., 2008). Despite using a substantially less extreme discrepancy between eye gaze and head direction, our results support these previous findings with the difference between central and eccentric fixation conditions as a proportion of eccentricity (−0.46° with a 12° discrepancy; 3.8%) broadly resembling other studies (−3.1° with a 45° discrepancy; 6.9%; (Lewald, 1998)).

**Figure S1.**
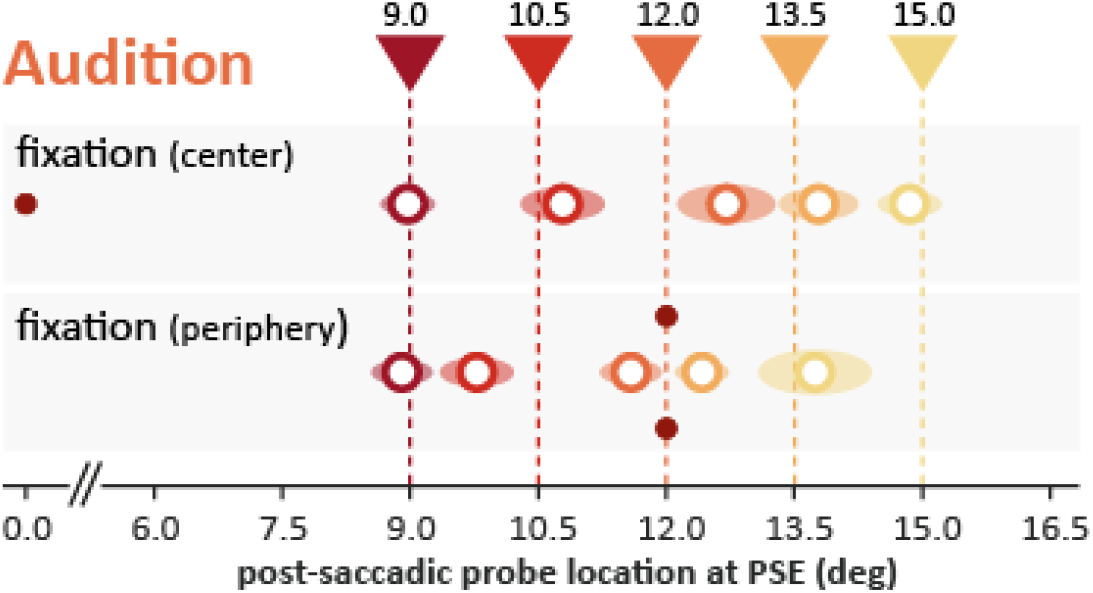
Auditory localization at fixation. Localization during central (top) and peripheral (bottom) fixation. Arrows indicate the position of the initial pre-saccadic stimulus while the white dots represent the mean location from which the post-saccadic stimulus had to originate in order to be perceived as occurring from the same location (shaded regions = ±1 SEM). Auditory localization was extremely accurate regardless of fixation, with no significant biases in either condition.

**Figure S2.**
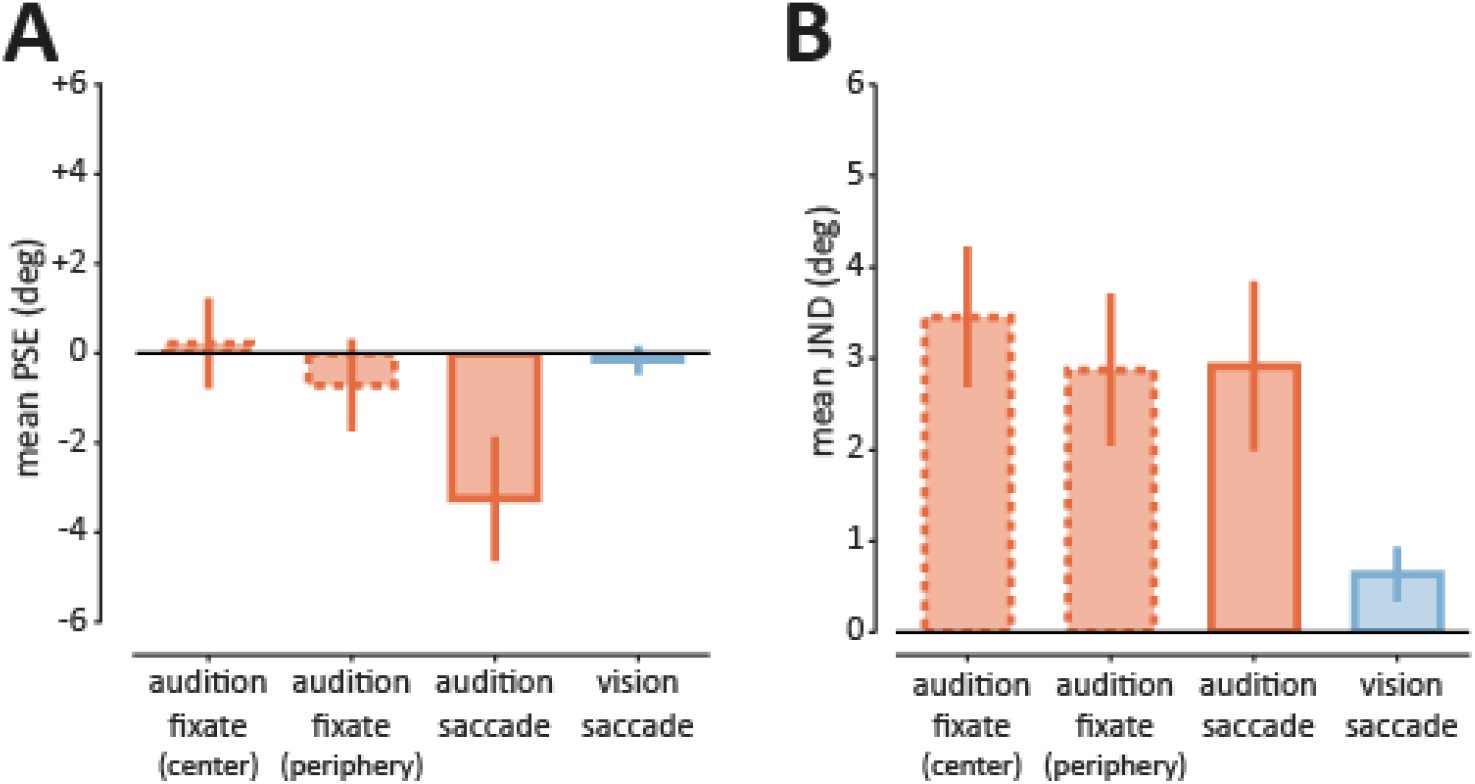
Static localization performance across modalities. **(A) Comparison of mean bias for fixation and static landmark conditions.** Mean change in the point of subjective equality (PSE) for the fixation tasks (dashed borders) and static landmark saccade tasks (full borders). There was a substantial bias for localizing auditory stimuli across the saccade, with negligible bias in the other conditions. **(B) Comparison of mean JND for fixation and static landmark conditions**. The mean just-noticeable-difference (JND) for the fixation tasks (dashed borders) and static landmark saccade tasks (full borders). While auditory localization was relatively precise in all conditions, peripheral viewing of the stimuli (due to peripheral fixation or due to an intervening saccade) resulted in a slight improvement. Unsurprisingly, visual localization was substantially more precise than auditory localization.

**Table S1.**
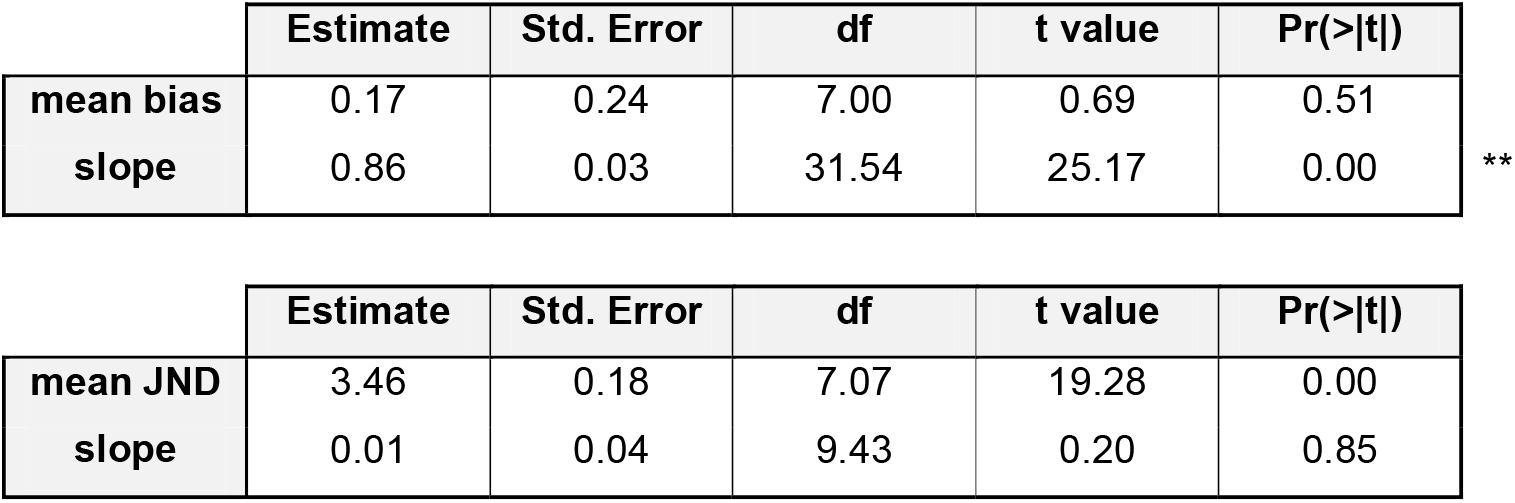
Auditory fixate (center)

**Table S2.** Auditory fixate (periphery)

|  | Estimate | Std. Error | df | t value | Pr(>\|t\|) |  |
| --- | --- | --- | --- | --- | --- | --- |
| **mean bias** | -0.75 | 0.21 | 7.02 | -3.64 | 0.01 | **\*\*** |
| **slope** | 0.76 | 0.04 | 7.67 | 17.08 | 0.00 | **\*\*** |
| **mean JND** | 3.00 | 0.23 | 7.00 | 13.07 | 0.00 | **\*\*** |
| **slope** | -0.01 | 0.04 | 7.00 | -0.20 | 0.85 |  |

**Table S3.**
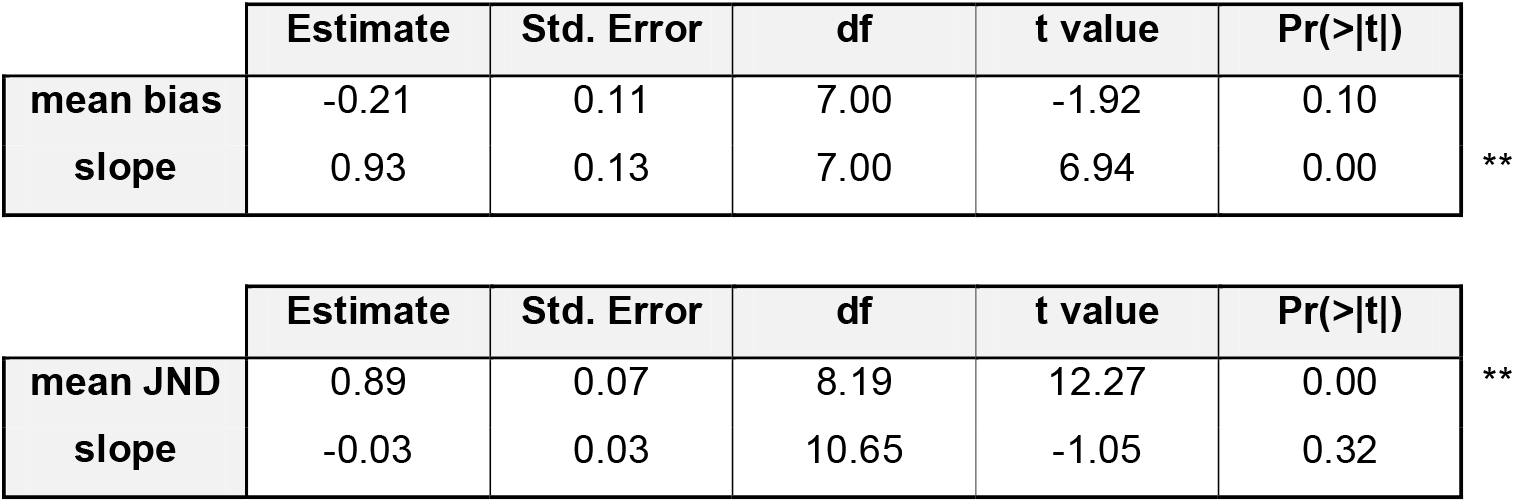
Visual saccade (no displacement)

**Table S4.** Auditory saccade (no displacement)

|  | Estimate | Std. Error | df | t value | Pr(>\|t\|) |  |
| --- | --- | --- | --- | --- | --- | --- |
| **mean bias** | -3.23 | 0.35 | 7.00 | -9.30 | 0.00 | **\*\*** |
| **slope** | 0.57 | 0.11 | 7.00 | 4.98 | 0.00 | **\*\*** |
| **mean JND** | 3.05 | 0.20 | 7.02 | 15.23 | 0.00 | **\*\*** |
| **slope** | -0.03 | 0.05 | 16.34 | -0.60 | 0.56 |  |

**Table S5.** Visual saccade (across displacements)

| Random effects | Name | Variance | Std.Dev. |
| --- | --- | --- | --- |
| Subject | (Intercept) | 0.023 | 0.152 |
|  | pre-saccadic location | 0.081 | 0.284 |
|  | landmark displacement | 0.007 | 0.081 |
|  | interaction | 0.000 | 0.009 |
| Residual |  | 0.381 | 0.617 |

| Fixed effects | Name | Estimate | Std. Error | df | t value | Pr(>\|t\|) |  |
| --- | --- | --- | --- | --- | --- | --- | --- |
| Subject | (Intercept) | -0.344 | 0.070 | 7.719 | -4.890 | 0.001 | ** |
|  | pre-saccadic location | 1.001 | 0.103 | 7.005 | 9.725 | 0.000 | ** |
|  | landmark displacement | 0.828 | 0.036 | 7.316 | 23.053 | 0.000 | ** |
|  | interaction | 0.005 | 0.011 | 39.718 | 0.417 | 0.679 |  |

**Table S6.** Auditory saccade (across displacements)

| Random effects | Name | Variance | Std.Dev. |
| --- | --- | --- | --- |
| Subject | (Intercept) | 0.862 | 0.928 |
|  | pre-saccadic location | 0.045 | 0.212 |
|  | landmark displacement | 0.014 | 0.120 |
|  | interaction | 0.000 | 0.017 |
| Residual |  | 0.829 | 0.910 |

| Fixed effects | Name | Estimate | Std. Error | df | t value | Pr(>\|t\|) |  |
| --- | --- | --- | --- | --- | --- | --- | --- |
| Subject | (Intercept) | -3.356 | 0.334 | 7.001 | -10.032 | 0.000 | ** |
|  | pre-saccadic location | 0.514 | 0.081 | 7.001 | 6.351 | 0.000 | ** |
|  | landmark displacement | 0.357 | 0.052 | 7.096 | 6.870 | 0.000 | ** |
|  | interaction | 0.009 | 0.015 | 22.261 | 0.596 | 0.557 |  |

## References

Aagten-Murphy D, Bays PM. 2018. Functions of Memory Across Saccadic Eye MovementsCurrent Topics in Behavioral Neurosciences. Springer, Berlin, Heidelberg. pp. 1–29.

Alais D, Burr D. 2004. The ventriloquist effect results from near-optimal bimodal integration. Curr Biol CB 14:257–262. doi:10.1016/j.cub.2004.01.029

Atsma J, Maij F, Koppen M, Irwin DE, Medendorp WP. 2016. Causal Inference for Spatial Constancy across Saccades. PLOS Comput Biol 12:e1004766. doi:10.1371/journal.pcbi.1004766

Bates D, Mächler M, Bolker B, Walker S. 2015. Fitting Linear Mixed-Effects Models Using lme4. J Stat Softw 67:1–48. doi:10.18637/jss.v067.i01

Batista AP, Buneo CA, Snyder LH, Andersen RA. 1999. Reach Plans in Eye-Centered Coordinates. Science 285:257–260. doi:10.1126/science.285.5425.257

Boon PJ, Belopolsky AV, Theeuwes J. 2016. The Role of the Oculomotor System in Updating Visual-Spatial Working Memory across Saccades. PLOS ONE 11:e0161829. doi:10.1371/journal.pone.0161829

Brainard DH. 1997. The Psychophysics Toolbox. Spat Vis 10:433–436.

Bridgeman B. 2007. Efference copy and its limitations. Comput Biol Med, Vision and Movement in Man and Machines 37:924–929. doi:10.1016/j.compbiomed.2006.07.001

Bridgeman B, Hendry D, Stark L. 1975. Failure to detect displacement of the visual world during saccadic eye movements. Vision Res 15:719–722. doi:10.1016/0042-6989(75)90290-4

Byrne PA, Crawford JD. 2010. Cue Reliability and a Landmark Stability Heuristic Determine Relative Weighting Between Egocentric and Allocentric Visual Information in Memory-Guided Reach. J Neurophysiol 103:3054–3069. doi:10.1152/jn.01008.2009

Camors D, Jouffrais C, Cottereau BR, Durand JB. 2015. Allocentric coding: Spatial range and combination rules. Vision Res 109:87–98. doi:10.1016/j.visres.2015.02.018

Campos M, Cherian A, Segraves MA. 2006. Effects of eye position upon activity of neurons in macaque superior colliculus. J Neurophysiol 95:505–526. doi:10.1152/jn.00639.2005

Cassanello CR, Ohl S, Rolfs M. 2016. Saccadic adaptation to a systematically varying disturbance. J Neurophysiol 116:336–350. doi:10.1152/jn.00206.2016

Cavanagh P, Hunt AR, Afraz A, Rolfs M. 2010. Visual stability based on remapping of attention pointers. Trends Cogn Sci 14:147–153. doi:10.1016/j.tics.2010.01.007

Cavanaugh J, Berman RA, Joiner WM, Wurtz RH. 2016. Saccadic Corollary Discharge Underlies Stable Visual Perception. J Neurosci 36:31–42. doi:10.1523/JNEUROSCI.2054-15.2016

Cohen YE, Andersen RA. 2002. A common reference frame for movement plans in the posterior parietal cortex. Nat Rev Neurosci 3:553–562. doi:10.1038/nrn873

Collins T. 2010. Extraretinal signal metrics in multiple-saccade sequences. J Vis 10:7–7. doi:10.1167/10.14.7

Collins T, Rolfs M, Deubel H, Cavanagh P. 2009. Post-saccadic location judgments reveal remapping of saccade targets to non-foveal locations. J Vis 9:29–29. doi:10.1167/9.5.29

Collins T, Wallman J. 2012. The relative importance of retinal error and prediction in saccadic adaptation. J Neurophysiol 107:3342–3348. doi:10.1152/jn.00746.2011

Cornelissen FW, Peters EM, Palmer J. 2002. The Eyelink Toolbox: Eye tracking with MATLAB and the Psychophysics Toolbox. Behav Res Methods Instrum Comput 34:613–617. doi:10.3758/BF03195489

Deubel H. 2004. Localization of targets across saccades: Role of landmark objects. Vis Cogn 11:173–202. doi:10.1080/13506280344000284

Deubel H, Bridgeman B, Schneider WX. 1998. Immediate post-saccadic information mediates space constancy. Vision Res 38:3147–3159.

Deubel H, Koch C, Bridgeman B. 2010. Landmarks facilitate visual space constancy across saccades and during fixation. Vision Res 50:249–259. doi:10.1016/j.visres.2009.09.020

Deubel H, Schneider WX, Bridgeman B. 1996. Postsaccadic target blanking prevents saccadic suppression of image displacement. Vision Res 36:985–996. doi:10.1016/0042-6989(95)00203-0

Duhamel JR, Colby CL, Goldberg ME. 1992. The updating of the representation of visual space in parietal cortex by intended eye movements. Science 255:90–92.

Engbert R, Mergenthaler K. 2006. Microsaccades are triggered by low retinal image slip. Proc Natl Acad Sci U S A 103:7192–7197. doi:10.1073/pnas.0509557103

Fiehler K, Wolf C, Klinghammer M, Blohm G. 2014. Integration of egocentric and allocentric information during memory-guided reaching to images of a natural environment. Front Hum Neurosci 8. doi:10.3389/fnhum.2014.00636

Ganmor E, Landy MS, Simoncelli EP. 2015. Near-optimal integration of orientation information across saccades. J Vis 15:8–8. doi:10.1167/15.16.8

Goossens HHLM, Opstal AJ van. 1999. Influence of Head Position on the Spatial Representation of Acoustic Targets. J Neurophysiol 81:2720–2736.

Groh JM, Trause AS, Underhill AM, Clark KR, Inati S. 2001. Eye position influences auditory responses in primate inferior colliculus. Neuron 29:509–518.

Grüsser O-J. 1983. Multimodal Structure of the Extrapersonal Space In: Hein A, Jeannerod M, editors. Spatially Oriented Behavior. Springer New York. pp. 327–352. doi:10.1007/978-1-4612-5488-1_18

Gruters KG, Murphy DLK, Jenson CD, Smith DW, Shera CA, Groh JM. 2018. The eardrums move when the eyes move: A multisensory effect on the mechanics of hearing. Proc Natl Acad Sci 201717948. doi:10.1073/pnas.1717948115

Hall NJ, Colby CL. 2011. Remapping for visual stability. Philos Trans R Soc Lond B Biol Sci 366:528–539. doi:10.1098/rstb.2010.0248

Harris LR, Smith AT. 2008. The coding of perceived eye position. Exp Brain Res 187:429–437. doi:10.1007/s00221-008-1313-0

Hartline PH, Vimal RLP, King AJ, Kurylo DD, Northmore DPM. 1995. Effects of eye position on auditory localization and neural representation of space in superior colliculus of cats. Exp Brain Res 104:402–408. doi:10.1007/BF00231975

Heed T, Buchholz VN, Engel AK, Röder B. 2015. Tactile remapping: from coordinate transformation to integration in sensorimotor processing. Trends Cogn Sci 19:251–258. doi:10.1016/j.tics.2015.03.001

Higgins E, Rayner K. 2015. Transsaccadic processing: stability, integration, and the potential role of remapping. Atten Percept Psychophys 77:3–27. doi:10.3758/s13414-014-0751-y

Jay MF, Sparks DL. 1987. Sensorimotor integration in the primate superior colliculus. I. Motor convergence. J Neurophysiol 57:22–34.

Jay MF, Sparks DL. 1984. Auditory receptive fields in primate superior colliculus shift with changes in eye position. Nature 309:345–347. doi:10.1038/309345a0

Karaminis T, Cicchini GM, Neil L, Cappagli G, Aagten-Murphy D, Burr D, Pellicano E. 2016. Central tendency effects in time interval reproduction in autism. Sci Rep 6. doi:10.1038/srep28570

Keating P, King AJ. 2015. Sound localization in a changing world. Curr Opin Neurobiol, Circuit plasticity and memory 35:35–43. doi:10.1016/j.conb.2015.06.005

King AJ. 2009. Visual influences on auditory spatial learning. Philos Trans R Soc Lond B Biol Sci 364:331–339. doi:10.1098/rstb.2008.0230

King AJ, Carlile S. 1993. Changes induced in the representation of auditory space in the superior colliculus by rearing ferrets with binocular eyelid suture. Exp Brain Res 94:444–455.

King AJ, Hutchings ME, Moore DR, Blakemore C. 1988. Developmental plasticity in the visual and auditory representations in the mammalian superior colliculus. Nature 332:73–76. doi:10.1038/332073a0

King AJ, Schnupp JWH, Doubell TP. 2001. The shape of ears to come: dynamic coding of auditory space. Trends Cogn Sci 5:261–270. doi:10.1016/S1364-6613(00)01660-0

Klingenhoefer S, Bremmer F. 2009. Perisaccadic localization of auditory stimuli. Exp Brain Res 198:411–423. doi:10.1007/s00221-009-1869-3

Knudsen EI, Knudsen PF. 1985. Vision guides the adjustment of auditory localization in young barn owls. Science 230:545–548.

Kopinska A, Harris LR. 2003. Spatial representation in body coordinates: evidence from errors in remembering positions of visual and auditory targets after active eye, head, and body movements. Can J Exp Psychol Rev Can Psychol Exp 57:23–37.

Krüger HM, Collins T, Englitz B, Cavanagh P. 2016. Saccades create similar mislocalizations in visual and auditory space. J Neurophysiol 115:2237–2245. doi:10.1152/jn.00853.2014

Kuznetsova A, Brockhoff PB, Christensen RHB. 2017. lmerTest Package: Tests in Linear Mixed Effects Models. J Stat Softw 82:1–26. doi:10.18637/jss.v082.i13

Lappe M, Awater H, Krekelberg B. 2000. Postsaccadic visual references generate presaccadic compression of space. Nature 403:892. doi:10.1038/35002588

Lee J, Groh JM. 2012. Auditory signals evolve from hybrid-to eye-centered coordinates in the primate superior colliculus. J Neurophysiol 108:227–242. doi:10.1152/jn.00706.2011

Lewald J. 1998. The effect of gaze eccentricity on perceived sound direction and its relation to visual localization. Hear Res 115:206–216. doi:10.1016/S0378-5955(97)00190-1

Lewald J, Ehrenstein WH. 1998. Auditory-visual spatial integration: a new psychophysical approach using laser pointing to acoustic targets. J Acoust Soc Am 104:1586–1597.

Lewald J, Ehrenstein WH. 1996. The effect of eye position on auditory lateralization. Exp Brain Res 108:473–485.

Lewald J, Guski R. 2003. Cross-modal perceptual integration of spatially and temporally disparate auditory and visual stimuli. Cogn Brain Res 16:468–478. doi:10.1016/S0926-6410(03)00074-0

Li J, Sajad A, Marino R, Yan X, Sun S, Wang H, Crawford JD. 2017. Effect of allocentric landmarks on primate gaze behavior in a cue conflict task. J Vis 17:20–20. doi:10.1167/17.5.20

Li W, Matin L. 1990. The influence of saccade length on the saccadic suppression of displacement detection. Percept Psychophys 48:453–458. doi:10.3758/BF03211589

McConkie GW, Currie CB. 1996. Visual stability across saccades while viewing complex pictures. J Exp Psychol Hum Percept Perform 22:563–581. doi:10.1037/0096-1523.22.3.563

Mullette-Gillman OA, Cohen YE, Groh JM. 2009. Motor-related signals in the intraparietal cortex encode locations in a hybrid, rather than eye-centered reference frame. Cereb Cortex N Y N 1991 19:1761–1775. doi:10.1093/cercor/bhn207

Mullette-Gillman OA, Cohen YE, Groh JM. 2005. Eye-centered, head-centered, and complex coding of visual and auditory targets in the intraparietal sulcus. J Neurophysiol 94:2331–2352. doi:10.1152/jn.00021.2005

Niemeier M, Crawford JD, Tweed DB. 2003. Optimal transsaccadic integration explains distorted spatial perception. Nature 422:76–80. doi:10.1038/nature01439

Opstal AJV, Hepp K, Suzuki Y, Henn V. 1995. Influence of eye position on activity in monkey superior colliculus. J Neurophysiol 74:1593–1610.

Ostendorf F, Dolan RJ. 2015. Integration of Retinal and Extraretinal Information across Eye Movements. PLOS ONE 10:e0116810. doi:10.1371/journal.pone.0116810

Paré M, Munoz DP. 2001. Expression of a re-centering bias in saccade regulation by superior colliculus neurons. Exp Brain Res 137:354–368. doi:10.1007/s002210000647

Parise CV, Harrar V, Spence C, Ernst M. 2011. Multisensory Integration: When Correlation Implies Causation. -Percept 2:901–901. doi:10.1068/ic901

Pavani F, Husain M, Driver J. 2008. Eye-movements intervening between two successive sounds disrupt comparisons of auditory location. Exp Brain Res 189:435–449. doi:10.1007/s00221-008-1440-7

Peck CK, Baro JA, Warder SM. 1995. Effects of eye position on saccadic eye movements and on the neuronal responses to auditory and visual stimuli in cat superior colliculus. Exp Brain Res 103:227–242.

Pelli DG. 1997. The VideoToolbox software for visual psychophysics: transforming numbers into movies. Spat Vis 10:437–442.

Poletti M, Burr DC, Rucci M. 2013. Optimal Multimodal Integration in Spatial Localization. J Neurosci 33:14259–14268. doi:10.1523/JNEUROSCI.0523-13.2013

Populin LC, Tollin DJ, Yin TCT. 2004. Effect of eye position on saccades and neuronal responses to acoustic stimuli in the superior colliculus of the behaving cat. J Neurophysiol 92:2151–2167. doi:10.1152/jn.00453.2004

Recanzone GH. 2009. Interactions of auditory and visual stimuli in space and time. Hear Res 258:89–99. doi:10.1016/j.heares.2009.04.009

Rolfs M. 2015. Attention in Active Vision: A Perspective on Perceptual Continuity Across Saccades. Perception 44:900–919. doi:10.1177/0301006615594965

Rolfs M, Szinte M. 2016. Remapping Attention Pointers: Linking Physiology and Behavior. Trends Cogn Sci 20:399–401. doi:10.1016/j.tics.2016.04.003

Rossetti Y, Tadary B, Prablanc C. 1994. Optimal contributions of head and eye positions to spatial accuracy in man tested by visually directed pointing. Exp Brain Res 97:487–496. doi:10.1007/BF00241543

Russo GS, Bruce CJ. 1994. Frontal eye field activity preceding aurally guided saccades. J Neurophysiol 71:1250–1253.

Schlack A, Sterbing-D’Angelo SJ, Hartung K, Hoffmann K-P, Bremmer F. 2005. Multisensory Space Representations in the Macaque Ventral Intraparietal Area. J Neurosci 25:4616–4625. doi:10.1523/JNEUROSCI.0455-05.2005

Schütz I, Henriques DYP, Fiehler K. 2013. Gaze-centered spatial updating in delayed reaching even in the presence of landmarks. Vision Res 87:46–52. doi:10.1016/j.visres.2013.06.001

Singmann H, Bolker B, Westfall J, Aust F, Højsgaard S, Fox J, Lawrence MA, Mertens U, Love J. 2016. afex: Analysis of factorial experiments. R package version 0.16–1.

Sommer MA, Wurtz RH. 2002. A Pathway in Primate Brain for Internal Monitoring of Movements. Science 296:1480–1482. doi:10.1126/science.1069590

Stricanne B, Andersen RA, Mazzoni P. 1996. Eye-centered, head-centered, and intermediate coding of remembered sound locations in area LIP. J Neurophysiol 76:2071–2076.

Szinte M, Cavanagh P. 2011. Spatiotopic apparent motion reveals local variations in space constancy. J Vis 11:4–4. doi:10.1167/11.2.4

Szinte M, Jonikaitis D, Rangelov D, Deubel H. 2018. Pre-saccadic remapping relies on dynamics of spatial attention. eLife 7. doi:10.7554/eLife.37598

Szinte M, Jonikaitis D, Rolfs M, Cavanagh P, Deubel H. 2016. Presaccadic motion integration between current and future retinotopic locations of attended objects. J Neurophysiol 116:1592–1602. doi:10.1152/jn.00171.2016

Vliegen J, Grootel TJV, Opstal AJV. 2004. Dynamic Sound Localization during Rapid Eye-Head Gaze Shifts. J Neurosci 24:9291–9302. doi:10.1523/JNEUROSCI.2671-04.2004

Wang X, Zhang M, Cohen IS, Goldberg ME. 2007. The proprioceptive representation of eye position in monkey primary somatosensory cortex. Nat Neurosci 10:640–646. doi:10.1038/nn1878

Watson WA. 1957. Contrast, Assimilation, and the Effect of Central Tendency. Am J Psychol 70:560–568. doi:10.2307/1419446

Weerts TC, Thurlow WR. 1971. The effects of eye position and expectation on sound localization. Percept Psychophys 9:35–39. doi:10.3758/BF03213025

Wexler M. 2003. Voluntary Head Movement and Allocentric Perception of Space. Psychol Sci 14:340–346. doi:10.1111/1467-9280.14491

Wexler M, Collins T. 2014. Orthogonal steps relieve saccadic suppression. J Vis 14:13–13. doi:10.1167/14.2.13

Wijdenes LO, Marshall L, Bays PM. 2015. Evidence for Optimal Integration of Visual Feature Representations across Saccades. J Neurosci 35:10146–10153. doi:10.1523/JNEUROSCI.1040-15.2015

Wolf C, Schütz AC. 2015. Trans-saccadic integration of peripheral and foveal feature information is close to optimal. J Vis 15:1–1. doi:10.1167/15.16.1

Wozny DR, Shams L. 2011. Recalibration of auditory space following milliseconds of crossmodal discrepancy. J Neurosci Off J Soc Neurosci 31:4607–4612. doi:10.1523/JNEUROSCI.6079-10.2011

Wurtz RH. 2008. Neuronal mechanisms of visual stability. Vision Res, Vision Research Reviews 48:2070–2089. doi:10.1016/j.visres.2008.03.021

Zella JC, Brugge JF, Schnupp JW. 2001. Passive eye displacement alters auditory spatial receptive fields of cat superior colliculus neurons. Nat Neurosci 4:1167–1169. doi:10.1038/nn773

Zimmermann E, Lappe M. 2010. Motor signals in visual localization. J Vis 10:2–2. doi:10.1167/10.6.2

Zwiers MP, Van Opstal AJ, Paige GD. 2003. Plasticity in human sound localization induced by compressed spatial vision. Nat Neurosci 6:175–181. doi:10.1038/nn999

Zwiers MP, Versnel H, Van Opstal AJ. 2004. Involvement of monkey inferior colliculus in spatial hearing. J Neurosci Off J Soc Neurosci 24:4145–4156. doi:10.1523/JNEUROSCI.0199-04.2004

